# High-level representations in human occipito-temporal cortex are indexed by distal connectivity

**DOI:** 10.1101/2021.02.22.432202

**Authors:** Jon Walbrin, Jorge Almeida

## Abstract

Human object recognition is dependent on occipito-temporal cortex, but a complete understanding of the complex functional architecture of this area must account for how it is connected to the wider brain. Converging functional magnetic resonance imaging evidence shows that univariate responses to different categories of information (e.g. faces, bodies, & non-human objects) are strongly related to, and potentially shaped by, functional and structural connectivity to the wider brain. However, to date, there have been no systematic attempts to determine how distal connectivity and complex local high-level responses in occipito-temporal cortex (i.e. multivoxel response patterns) are related. Here, we show that distal functional connectivity is related to, and can reliably index, high-level representations for several visual categories (i.e. tools, faces, & places) within occipito-temporal cortex; that is, voxels sets that are strongly connected to distal brain areas show higher pattern discriminability than less well-connected sets do. We further show that, in several cases, pattern discriminability is higher in sets of well-connected voxels than sets defined by ‘local’ activation (e.g. strong amplitude responses to faces in fusiform face area). Together, these findings demonstrate the important relationship between the complex functional organization of occipito-temporal cortex and wider brain connectivity.

**Significance statement:** Human object recognition relies strongly on occipito-temporal cortex (OTC), yet responses in this broad area are often considered in relative isolation to the rest of the brain. We employ a novel ‘connectivity-guided’ voxel selection approach with functional MRI data to show higher sensitivity to information (i.e. higher multivoxel pattern discriminability) in voxel sets that share strong connectivity to distal brain areas, relative to: 1) voxel sets that are less-strongly connected; and in several cases 2) voxel sets that are defined by strong ‘local’ response amplitude. These findings underscore the importance of distal contributions to local processing in OTC.

## Introduction

Human object recognition is a rapid process that relies heavily on occipito-temporal cortex (OTC; e.g. Grill-Spector & Malach, 2004), and there have been extensive efforts to fully characterize the complex functional organization of this area (e.g. Grill-Spector & Weiner, 2014; Op de Beeck et al., 2019; Peelen & Downing, 2017). Convergent functional magnetic resonance imaging (fMRI) findings show coarse-grain organization of OTC as evidenced by spatially clustered ‘category-preferring’ responses, that is, regions that show enhanced fMRI response amplitude for one category over others (e.g. faces, tools, & places/scenes: Almeida et al., 2013; Beauchamp & Martin, 2007; Chao & Martin, 2000; Downing et al., 2006; Epstein and Kanwisher, 1998; Kanwisher et al., 1997; Kristensen et al., 2016), along with finer-grain organization via ‘patchy’ organization of OTC (i.e. sparsely distributed cortical patches that respond strongly to different information; e.g. Grill-Spector et al., 2006; Weiner & Grill-Spector, 2010) that are well-captured with multivoxel pattern analysis (MVPA) techniques (Haxby et al., 2001; Kamitani & Tong, 2005).

However, a complete understanding of the functional architecture of OTC must account for how this broad area interfaces with the wider brain. Indeed, connectivity is a major constraint on the functional organization of cerebral cortex in general, such that the functional response of a given region is partially determined by the integration of relevant information shared via structural and functional connectivity to other brain regions (e.g. Garcea et al., 2019; Lee et al., 2019; Mahon & Caramazza, 2011; Sporns & Zwi, 2004; Sporns, 2004). More specifically, category-preferring OTC responses are functionally coupled with, and modulated by, distal regions that share the same category-preference (e.g. tool responses in medial fusiform gyrus are shaped by inferior parietal cortex; Amaral et al., under review; Chen et al., 2017; Garcea et al., 2019; Lee et al., 2019); similarly, OTC responses for multiple visual categories (e.g. faces, objects, bodies, & places) can be reliably predicted from patterns of white matter connectivity to the wider brain (Osher et al., 2016; Saygin et al., 2012; Saygin et al., 2016).

The preceding evidence demonstrates a clear relationship between distal connectivity and functional local OTC responses at the level of individual voxels. However, the extent to which connectivity relates to complex distributed functional responses (i.e. multivoxel pattern decoding), is not yet understood. Here, we show that the discriminability of distributed multivoxel response patterns in OTC is related to – and importantly, can be *indexed* by – patterns of distal connectivity; that is, sets of voxels that afford high pattern discriminability of different object categories can be identified by the strength of connectivity they share with distal brain areas. Specifically, our results demonstrate that: 1) ‘Most-connected’ grey matter voxel sets consistently yield higher pattern discriminability than ‘least-connected’ sets do, and; 2) Most-connected voxel sets are partially distinct from – and, in several cases, afford significantly higher pattern discriminability than – ‘most-activated’ voxel sets do (i.e. sets defined by strongest amplitude responses). In summary, these findings demonstrate a compelling relationship between distal connectivity and locally distributed functional responses in OTC.

## Materials and Methods

### Participants

20 right-handed undergraduate adult participants (mean age: 22.1 years; SD: 5.4; 14 females) gave informed consent and were reimbursed with university course credit. Head motion was not excessive for any subject (i.e. no >2mm scan-to-scan spikes), so all data was used. Ethical procedures were approved by the Faculty of Psychology and Educational Sciences of the University of Coimbra ethics board.

### MRI scanning parameters

Scanning was performed with a Siemens Tim Trio 3T MRI scanner (Siemens Healthineers, Erlangen, Germany) with a 12-channel head coil at the University of Coimbra. Functional images were acquired with the following parameters: T2*-weighted single-shot echo-planar imaging pulse sequence, TR = 2000ms, TE = 30ms, flip angle = 90°, 40 interleaved axial slices (no gap), acquisition matrix = 96 x 96 with field of view = 256mm, with a voxel size of 2.3 x 2.3 x 3mm. Structural T1-weighted images were obtained using an MPRAGE (magnetization prepared rapid gradient echo) sequence with the following parameters: TR = 2530ms, TE = 3.29ms, in 1.7ms steps, total acquisition time = 136s, FA = 8°, acquisition matrix = 256 × 256, with field of view 256mm, and voxel size = 1 mm^3^.

### Task

Participants completed 6 runs of a blocked-design task, where they centrally fixated grey-scaled images (400 x 400 pixels; ∼10° of visual angle) of tools, faces, and places (animal images, as well as phase-scrambled variants of these categories were also presented, but were not analysed here). Each run consisted of alternating 6s blocks of stimuli and 6s fixation, with 16s fixation at the beginning and end of each run (run length: 176s = 88 TRs); 2 blocks were presented for each of the categories (and 1 block for each of the phase-scrambled conditions). Block order was randomized across runs.

### Pre-processing

Pre-processing was performed with SPM12 (fil.ion.ucl.ac.uk/spm/software/spm12). This entailed slice-timing correction, re-alignment (and re-slicing), co-registration, and segmentation. Segmented grey matter maps were co-registered and warped to subject’s functional image space for later masking out white matter voxels. A duplicate set of functional data was normalized and smoothed for the sole purpose of identifying group-level activation peaks for creating ‘search spaces’ for each target area. All default SPM12 parameters were used (except for normalized data where output the voxel size was 3mm^3^ and a 6mm^3^ FWHM Gaussian smoothing kernel was used).

General linear model estimation was performed in SPM12 and all analyses were performed in subject space. Block durations and onsets for each experimental condition were modelled by convolving the corresponding box-car time-course with a canonical hemodynamic response function (without time or dispersion derivatives), with a high-pass filter of 256s and autoregressive AR(1) model. Beta maps were generated on a run-wise basis, yielding one regressor per condition, along with 6 rigid-motion regressors (and an intercept regressor). T-maps were estimated for the contrasts described below.

The pre-processed functional data was duplicated and de-noising was performed with the CONN Toolbox (Whitfield-Gabrieli & Nieto-Castanon, 2012) by regressing out task-related effects (i.e. haemodynamic response convolved with blocks for each condition), along with other head motion (6 rigid-motion regressors + 6 first-order temporal derivatives) and physiological noise related variables (mean global signal estimated from all white matter and cerebrospinal fluid voxels, along with outlier scan removal), and band-pass filtered (0.01-0.1hz). Previous work shows that this approach successfully removes task-related signal, resulting in time-course data that this very similar to resting-state fMRI signal (e.g. Fair et al., 2007).

### Voxel selection

We employed a ‘connectivity-guided voxel selection’ approach in 6 target regions: Tool-preferring medial fusiform gyrus (MFus) and posterior medial temporal gyrus (PMTG); face-preferring fusiform face area (FFA) and occipital face area (OFA); and place-preferring parahippocampal place area (PPA) and occipital place area (OPA). The following regions outside of OTC were also used for connectivity seeding: Tool-preferring inferior parietal lobe (IPL) and superior parietal lobe (SPL); face-preferring superior temporal sulcus (STS-F), and; place-preferring retrosplenial cortex (RSC). Thus, voxel selection within a given target region depended on connectivity to all other regions (both within and outside of OTC) that shared the same category-preference (e.g. PMTG, IPL, & SPL served as ‘seed’ regions for voxel selection in MFus). Analyses were restricted to left-hemisphere tool regions, and right hemisphere face and place regions, based on widely-observed hemispheric asymmetries (e.g. Downing et al., 2006); however, we also observed the same pattern of results in the ‘opposite’ hemisphere, for each set of regions.

Target region masks (i.e. ‘search spaces’ for voxel selection) were created by centring a 15mm-radius sphere at the most-activated voxel (uncorrected p<.05), based on group-level activation in normalised space. For tool-, face-, and place-preferring regions, activation was based on the following t-contrasts respectively: Tools > [faces + places + animals]; faces > [tools + places + animals]; and places > [tools + faces + animals]. Voxels that overlapped between two adjacent search spaces for the same category (e.g. OFA and FFA) were removed; search spaces between categories (e.g. tool-preferring MFus and face-preferring FFA) were free to overlap. Target region masks were then inverse-registered to each subject’s own brain space, and white-matter (and cerebellum) voxels were removed. All target region masks contained >300 grey-matter voxels. ‘Seed’ regions were then defined as the 100 most-activated voxels (for the same contrasts described above, based on each subject’s own activity (i.e. t-values)) within each target region.

For each target region, a functional connectivity matrix was calculated that described the time-course correlation (Fisher-transformed Pearson’s *r* coefficient) between each voxel and *all other same-category seed voxels* (e.g. 350 MFus target regions voxels x 100 PMTG + 100 IPL + 100 SPL seed region voxels). The mean correlation for each target region voxel (across all seed regions) was then obtained, and the 100 most highly connected and 100 least highly connected target region voxels were selected, respectively.

We also compared sets of most-connected voxels with sets of 100 most highly activated voxels based on the corresponding t-contrast for each decoding analysis (e.g. tools > faces t-values were used for tools vs. faces decoding). Importantly, due to potential circularity problems (Kriegeskorte et al., 2009; e.g. exaggerated tools vs. faces decoding accuracy might result if voxel sets are defined with the exact same data) data was independently split for voxel selection and decoding, as follows. Subject data (both task & task-regressed connectivity datasets) was divided into 3 splits (2 runs each). A leave-one-split-out approach was adopted (for generating and testing both connectivity and activity voxel sets) where one split of data was used for voxel selection while the remaining 2 splits were used in the corresponding decoding fold (iterated 3 times, so that each split was used for voxel selection). Within each target region, voxels did not overlap for the most-connected- and least-connected sets (but overlap was unconstrained between most-connected- and most-activated voxel sets).

### Signal-to-noise-ratio analysis

To test whether subtle differences in signal-to-noise-ratio (SNR) might explain a potential decoding advantage in most-connected-relative to least-connected voxel sets – that is, higher SNR in most-connected voxel sets might partially account for higher decoding compared to least-connected voxel sets – we directly compared SNR between voxel sets as follows.

Whole brain maps that describe the voxel-wise temporal SNR (i.e. mean signal amplitude / SD; e.g. Triantafyllou et al., 2005) for each run of task data, were generated, for each subject. Mean SNR values for most-connected- and least-connected voxel sets were then obtained for each subject across all 6 runs of data (within each target area, and averaged across voxel sets from all 3 data splits), and entered into 2-way ANOVA (voxel selection type x region). These analyses revealed an effect in the opposite direction – that is SNR was slightly higher in least-connected-than most-connected voxel sets (main effect of voxel selection: (F(1,19) = 25.51, p < .001, η_p_^2^ = .573; most-connected > least connected (post-hoc contrast): t(19) = −5.05, p < .001), indicating that any potential decoding advantage in most-connected voxel sets relative to least-connected voxel sets, is not attributable to higher SNR in most-connected voxel sets.

### Multivoxel pattern decoding

Decoding was implemented with the CoSMoMVPA Toolbox (Oosterhof et al., 2016). A split-half Pearson’s *r* correlation decoding approach was used (e.g. see Haxby et al., 2001) as a measure of discriminability between each relevant pair of conditions. This is a powerful decoding approach that performs equivalently to commonly-used linear classifiers (Misaki et al., 2010).

Decoding was performed across 3 decoding folds (i.e. voxels selected with 1 data split and decoding performed with the 2 left-out data splits, with a different data split used for voxel selection for each decoding fold). 2 decoding comparisons were run for each category to ensure the generalizability of effects (i.e. tool-preferring regions: tools vs. faces & tools vs. places; face-preferring regions: faces vs. tools & faces vs. places; place-preferring regions: places vs. tools & places vs. faces). For each decoding comparison pair (e.g. tools vs. faces), patterns for each condition were correlated across the 2 designated decoding splits of data (i.e. 2 runs per split), yielding a 2 (split) x 2 (category) confusion matrix, where the mean between-category correlation (off-diagonal cells) was subtracted from the mean within-category correlation (on-diagonal cells); thus, a positive decoding accuracy denotes greater within-category than between-category decoding (e.g. [tools-to-tools correlation + faces-to-faces correlation] > tools-to-faces correlations, across splits). Subjects’ decoding accuracy values were mean-averaged across decoding folds and entered into 3-way repeated measures ANOVAs (i.e. voxel selection type x region x decoding comparison), for each set of category-preferring regions, separately.

For conciseness, only ANOVA terms involving the factor ‘voxel selection type’ are reported here; specifically, we only report these effects at the highest descriptive level (i.e. for significant interactions involving voxel selection type, we report the corresponding post-hoc test; in the absence of a significant interaction term, we report the main effect of voxel selection type). A Bonferroni-corrected threshold was calculated for each set of post-hoc t-tests (two-tailed) involving voxel selection type, and all reported tests survive correction unless otherwise stated.

### Matched-activation analyses

To ensure that any differences between most-connected- and least-connected voxel sets were not confounded by local activation to category information (e.g. differences between the two voxel sets in FFA might result from differences in mean activation differences to faces), a series of ‘matched-activation’ analyses were performed. This entailed selecting strongly-connected- and weakly-connected voxel sets under the constrained that they did not statistically differ by their average activation (t-values; e.g. for face regions, t-values were matched for each corresponding decoding analysis (e.g. faces > tools t-values were used for faces vs. tools decoding). This was achieved with a permutation approach as follows.

Voxels in each target region were median-split by their mean connectivity values (i.e. mean connectivity-correlation value to all seed voxels). 2 random subsets of 100 voxels – one each from the highest- and lowest half-splits – were then drawn and compared to ensure that their average activation values – *voxel t-values* – did not differ when compared via an independent t-test. 10,000 subset comparisons were performed but, crucially, only subset-pairs with *non-significant independent t-test statistics* were retained. Decoding was then performed with these ‘non-differing activation’ voxel sets, and averaged to create stable decoding estimates for the strongly-connected- and weakly-connected voxel sets, respectively.

This analysis was repeated across statistical 3 thresholds, retaining voxel set pairs that did not differ: 1) At a liberal threshold (i.e. two tailed independent t-test p-values > .10); 2) At an intermediate threshold (i.e. independent t-test statistics within the range t = +0.5 to −0.5), and; 3) At a strict threshold, where voxel set pairs were only accepted when the average activation was *lower* in strongly-connected sets (i.e. independent t-test statistics within the negative range of t = 0 to −0.5).

We initially ran 3-way repeated measures ANOVAs (voxel selection x decoding comparison x region) for tools, faces, and places separately, as in the main analyses to test these results. However, we observed reduced degrees of freedom for these analyses, indicating that these constraints were not always met in all subjects and regions (e.g. connectivity and activity were less independent of each other for some region in some subjects, such that mean activation always differed between voxel sets). To preserve statistical power, we ran follow-up 2-way repeated measures ANOVAs (voxel selection x decoding comparison) for each region separately, if the initial 3-way ANOVA indicated that at least 3 subjects did not meet this constraint; as such, if the constraint was met in one region but not the other in a given subject, this would allow for their data to be retained when testing the ‘surviving’ region. Specifically, 3-way ANOVA results are reported for face regions as only 1 subject failed to meet this constraint (across all 3 matched-activation analysis thresholds). Due to higher subject drop-out for the other 3-way ANOVAs (i.e. for tools & places, respectively), region-wise 2-way ANOVA results are reported for the 2 more conservative thresholds (but 3-way ANOVA results are reported at the most liberal threshold, where only 2 subjects failed to meet this constraint). The number of remaining subjects per analysis is reported in the results section.

### ‘Good seed’ searchlight analysis

To complement the main analyses that used a-priori seed regions, we ran a searchlight analysis to determine which regions across the entire brain constituted ‘good seeds’ (i.e. regions with connectivity that yields higher decoding in most-connected-, compared to least-connected target region voxels). For each target region (e.g. MFus), a searchlight consisting of approximately 100 contiguous voxels was centred on each given grey matter voxel of the brain (excluding the given target region), and the mean time-course for those voxels was correlated with the target region for voxel selection. Decoding was performed and accuracy values were then assigned to the central voxel of the corresponding searchlight. This was performed for each subject, across all analysis variants (i.e. for each category, 2 target regions x 2 binary decoding comparisons x 2 voxel sets (i.e. most- & least-connected)).

The same decoding approach as in the main analyses (i.e. with a-priori seeds) was adopted here, except that decoding was performed with all 6 runs of data in a *single decoding fold* (i.e. where run-averaged patterns between the 3 odd and 3 even runs were correlated), rather than adhering to the data split scheme imposed in the previous analyses (i.e. 3 decoding folds). This was done for two reasons: 1) Because activation was not used for comparative voxel selection here, data circularity problems do not apply; and 2) this demonstrates the generalizability of the distal connectivity decoding effect with a different decoding scheme (we also ran these analyses with the same split-scheme as in the main analyses and observed virtually identical results).

For group-level inference, paired t-tests with threshold-free cluster enhancement (TFCE; see Smith & Nichols, 2009) based on 10,000 Monte-Carlo simulations were run with subjects’ most-connected- and least-connected voxel selection searchlight maps (maps were normalised and smoothed with a 6mm FWHM kernel, beforehand). The resulting group-level maps were thresholded at Z > 1.65 and projected to a surface rendered brain in SPM12 for visualization. In short, these maps show regions that constitute ‘good seeds’, yielding a significant decoding effect (i.e. seeding from these regions results in higher decoding for the most-connected-than least-connected voxel sets in the corresponding target region).

## Results

Across all 6 target regions, higher decoding accuracy was observed for most-connected-than least-connected voxel sets (see fig. 1, upper row bars). This effect was shown for both tool regions (i.e. MFus & PMTG; main effect of voxel selection type: F(1,19) = 46.85, p < .001, η_p_^2^ = .711), both face regions (i.e. FFA & OFA; voxel selection type x decoding comparison (interaction): F(1,19) = 14.64, p < .001, η_p_^2^ = .435; faces vs. places (post-hoc): t(24.30) = 5.79, p < .001; faces vs. tools (post-hoc): t(24.30) = 3.09, p = .005), and both place regions (i.e. PPA & OPA; voxel selection type x decoding comparison (interaction): F(1,19) = 12.11, p = .003, η_p_^2^ = .389; places vs. faces (post-hoc): t(28.32) = 7.69, p < .001; places vs. tools (post-hoc): t(28.32) = 4.52, p < .001). Thus, decoding accuracy is consistently higher in voxel sets that are most-rather than least-distally connected.

**Figure 1.**
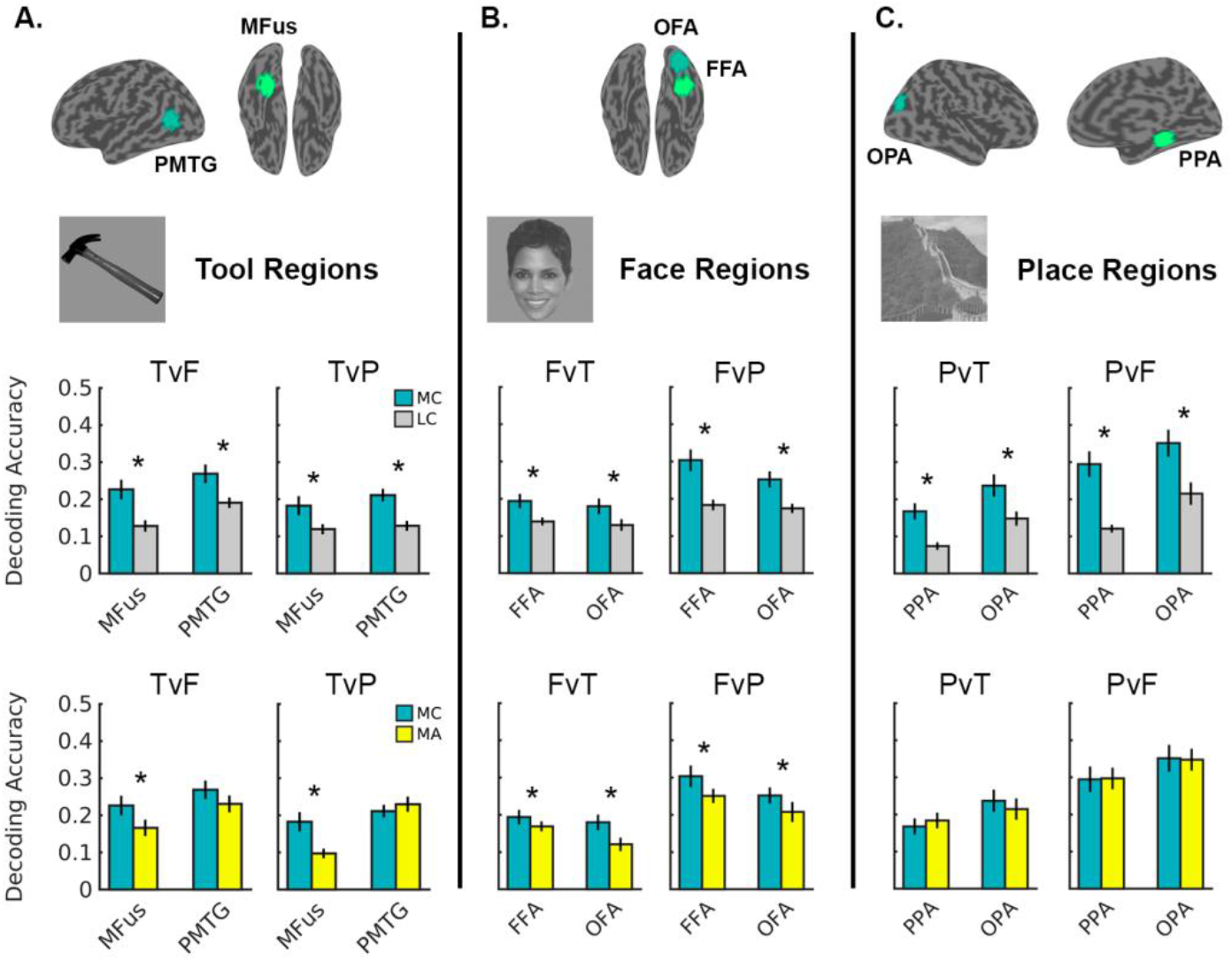
Mean decoding accuracy for most-connected- (MC), least-connected- (LC), & most-activated (MA) voxel sets, for (a) tool, (b) face, & (c) place regions. Upper row bar charts: MC vs. LC decoding. Lower row bar charts: MC vs. MA decoding. Tool regions: MFus = medial fusiform gyrus; PMTG = posterior middle temporal gyrus. Face regions: FFA = fusiform face area; OFA = occipital face area. Place regions: PPA = parahippocampal place area; OPA = occipital place area. Decoding comparisons: Tools vs. faces (TvF); tools vs. places (TvP); faces vs. tools (FvT); faces vs. places (FvP); places vs. tools (PvT); places vs. faces (PvF). * = significant MC > LC effect (Bonferroni-corrected p < .05). Error bars are SEM. Target region search spaces are shown on a surface brain, along with example stimuli in the upper portion of the figure.

Previous evidence shows that distal functional connectivity is correlated with task-based activation in OTC (e.g., Chen et al., 2017; Amaral et al., under review). It is therefore possible that most-connected voxels are effectively the same as those showing the strongest local activation, and therefore might yield equivalent decoding performance. We tested this by comparing decoding accuracy in most-connected voxel sets with those that were most-activated by their ‘preferred’ stimulus category (both sets were free to overlap). Interestingly, decoding accuracy was never statistically lower in most-connected-relative to most-activated voxel sets; indeed, decoding in most-connected voxel sets was almost always equal or higher than decoding in most-activated voxel sets (see fig. 1, lower row bars). For tool-preferring regions, decoding accuracy was significantly higher for most-connected-, relative to most-activated voxel sets (voxel selection x decoding comparison x region interaction: F(1,19) = 5.04, p = .037, η_p_^2^ = .210) in MFus (tools vs. faces (post-hoc): t(74.31) = 3.16, p = .002; tools vs. places (post-hoc): t(74.31) = 4.45, p < .001), but this trend was not significant in PMTG (tools vs. faces (post-hoc): t(74.31) = 1.99, p = .050; tools vs. places (post-hoc): t(74.31) = −0.97, p = .335). By contrast, this effect was significant in both FFA and OFA (main effect of voxel selection type: F(1,19) = 10.62, p = .004, η_p_^2^ = .358), but was not significant in either place region (all voxel selection ANOVA terms: p > .115).

These results show that decoding performance is not equivalent in most-connected- and most-activated voxel sets in several areas. Nevertheless, we further sought to test the relative independence of the effects observed in the original analysis by looking at whether the decoding differences between most-connected and least-connected voxel sets remain when potential differences in average activation between sets are controlled; that is, does greater decoding in most-connected-than least-connected voxel sets remain when average activation (t-values) between the two sets is closely controlled?

To test this, we ran a ‘matched activation’ permutation analysis where we median split each target area by voxel connectivity values, and randomly drew subsets of 100 strongly-connected- and 100 weakly-connected voxels but, crucially, only compared decoding performance in sets that did not statistically differ by their average activation (i.e. t-values; see methods for full details). In the first variant of this analysis, we retained pairs of voxel sets that did not statistically differ at a relatively liberal threshold (i.e. two-tailed p>.10) when running an independent t-test between the 2 voxel sets’ activation values (t-values)).

As before, decoding accuracy was significantly higher for most-connected-than least-connected voxel sets (see fig 2; upper row bars) in both tool regions (main effect of voxel selection: F(1,17) = 52.04, p < .001, η_p_^2^ = .754), both face regions (voxel selection type x decoding comparison (interaction): F(1,19) = 23.22, p < .001, η_p_^2^ = .550; faces vs. places (post-hoc): t(23.57) = 6.60, p < .001; faces vs. tools (post-hoc): t(23.57) = 3.43, p = .002) and both place regions (voxel selection type x decoding comparison (interaction): F(1,17) = 10.39, p = .005, η_p_^2^ = .379; places vs. faces (post-hoc): t(25.9) = 7.81, p < .001; places vs. tools (post-hoc): t(25.9) = 4.79, p < .001).

**Figure 2.**
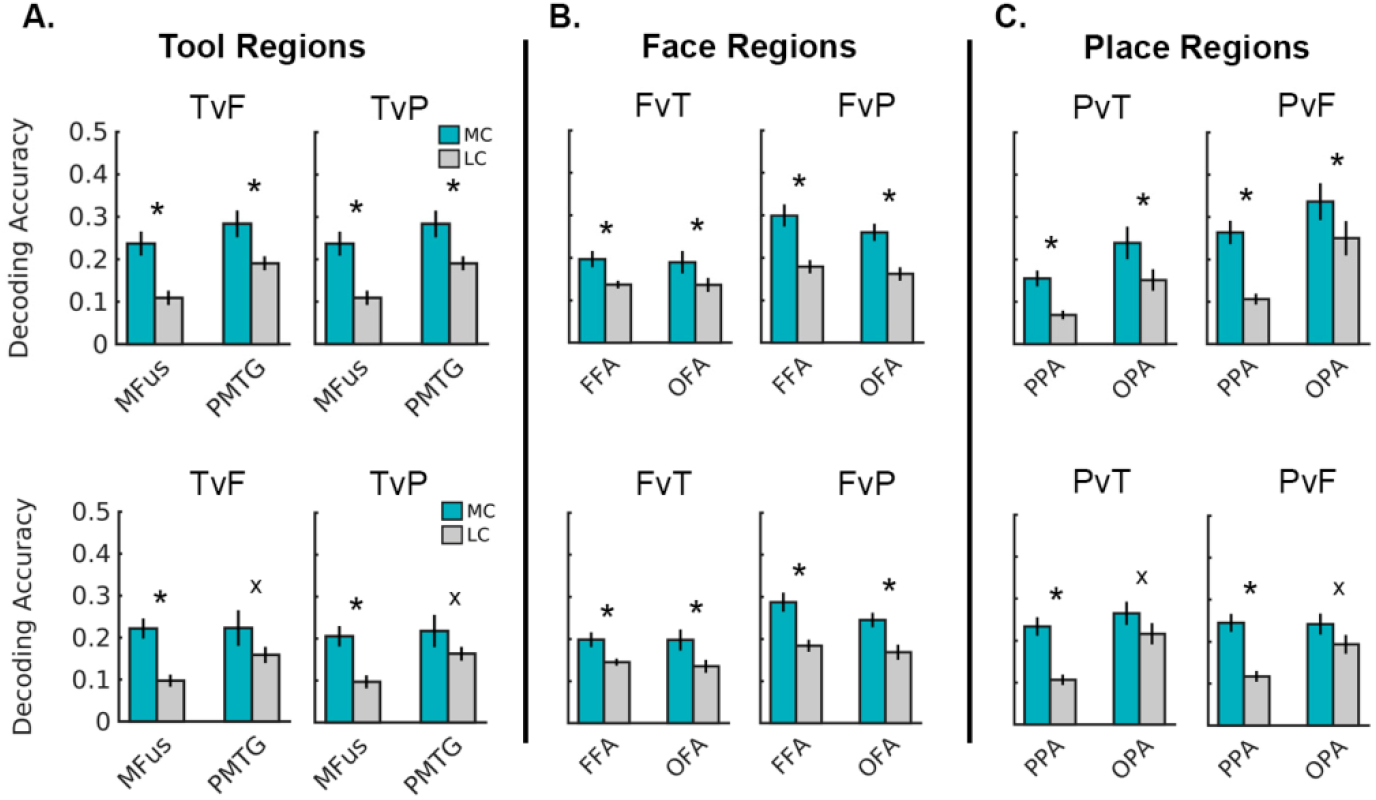
Mean decoding accuracy for most-connected- (MC) & least-connected (LC) voxel sets for ‘matched activation’ analyses: (a) tool-, (b) face-, & (c) place regions. Upper row bars: Liberal ‘non-different activation’ threshold (p>.10). Lower row bars: Strictest ‘non-different activation’ threshold (negative t-values between 0 to −0.5). Tool regions: MFus = medial fusiform gyrus; PMTG = posterior middle temporal gyrus. Face regions: FFA = fusiform face area; OFA = occipital face area. Place regions: PPA = parahippocampal place area; OPA = occipital place area. Decoding comparisons: Tools vs. faces (TvF); tools vs. places (TvP); faces vs. tools (FvT); faces vs. places (FvP); places vs. tools (PvT); places vs. faces (PvF). * = significant effect (Bonferroni-corrected p < .05). x = trend non-significant/underpowered analyses (PMTG N=8; OPA N=13). Error bars are SEM.

We next repeated this analysis under 2 stricter thresholds by only retaining voxel set pairs where: 1) Average activation was more closely matched between the 2 sets (i.e. independent t-tests that yielded t-statistics with the range of +0.5 to −0.5), and; 2) average activation was *lower* in most-connected voxel sets (i.e. independent t-tests that yielded negative t-statistics with the range of 0 to −0.5). Given these conservative criteria, we anticipated that these constraints would not be met in all subjects, and therefore the number of ‘surviving’ subjects are reported for each analysis.

Under the intermediate threshold (i.e. t-statistics between +0.5 to −0.5), higher decoding accuracy was observed for most-connected-than least-connected voxel sets in all regions. For MFus, FFA, OFA, and, PPA, 19 out of 20 subjects met this constraint (MFus: F(1,18) = 19.30, p < .001, η_p_^2^ = .517; FFA & OFA: F(1,18) = 23.59, p < .001, η_p_^2^ = .567; PPA: F(1,18) = 43.46, p < .001, η_p_^2^ = .707). These effects were also significant in PMTG and OPA where 13 and 14 subjects remained, respectively (PMTG: F(1,12) = 6.97, p = .022, η_p_^2^ = .367; OPA: F(1,13) = 4.83, p = .047, η_p_^2^ =.271).

Under the strictest constraint (i.e. t-statistics between 0 to −0.5), higher decoding accuracy was (again) observed for most-connected-than least-connected voxel sets across all regions (see fig 2; lower row bars). This trend was statistically significant in all regions where this constraint was met for at least 17 out of 20 subjects: MFus, FFA, OFA, and PPA (MFus (1,18) = 18.66, p < .001, η_p_^2^ = .509; FFA & OFA (post-hoc test; faces vs. tools): t(24.97) = 3.26, p = .003; FFA & OFA (post-hoc test; faces vs. places): t(24.97) = 5.04, p < .001; PPA: F(1,16) = 35.00, p < .001, η_p_^2^ = .686). In PMTG and OPA, these analyses were underpowered (i.e. only 8 and 13 subjects remained, respectively) and did not reach significance (PMTG: F(1,7) = 1.51, p = .258, η_p_^2^ = .178; OPA: F(1,12) = 2.81, p = .120, η_p_^2^ = .190). Although these trends are evident in figure 2 (lower row), these results show a lesser degree of independence between connectivity and activity measures in PMTG and OPA than the other regions.

Taken together, these analyses show strong decoding performance in highly-connected voxel sets; importantly, these distally well-connected voxel sets demonstrate a degree of independence from – and therefore, are not merely confounded by – local voxel activation (t-values).

Finally, we implemented whole-brain searchlight analyses (see fig. 3) for each target region; these analyses revealed regions (beyond a-priori ‘seed’ regions used in the preceding analyses) that afford ‘good seeding’ (i.e. regions with distal connectivity that yields higher decoding in most-connected-rather than least-connected voxel sets). Diffuse patterns of strong seeding in the wider brain were shown for all target regions. Notably, good seeding was observed in bilateral posterior temporal cortex (coincident with category-preferring OTC regions) and early visual cortex, as well as dorsal attention and task-general cognitive control regions (e.g. anterior inferior parietal sulcus, frontal eye fields, and pre-central gyrus) and this coverage comparable with previously observed functional connectivity patterns between OTC and the wider brain (Hutchison et al., 2014; Vogel et al., 2012).

**Figure 3.**
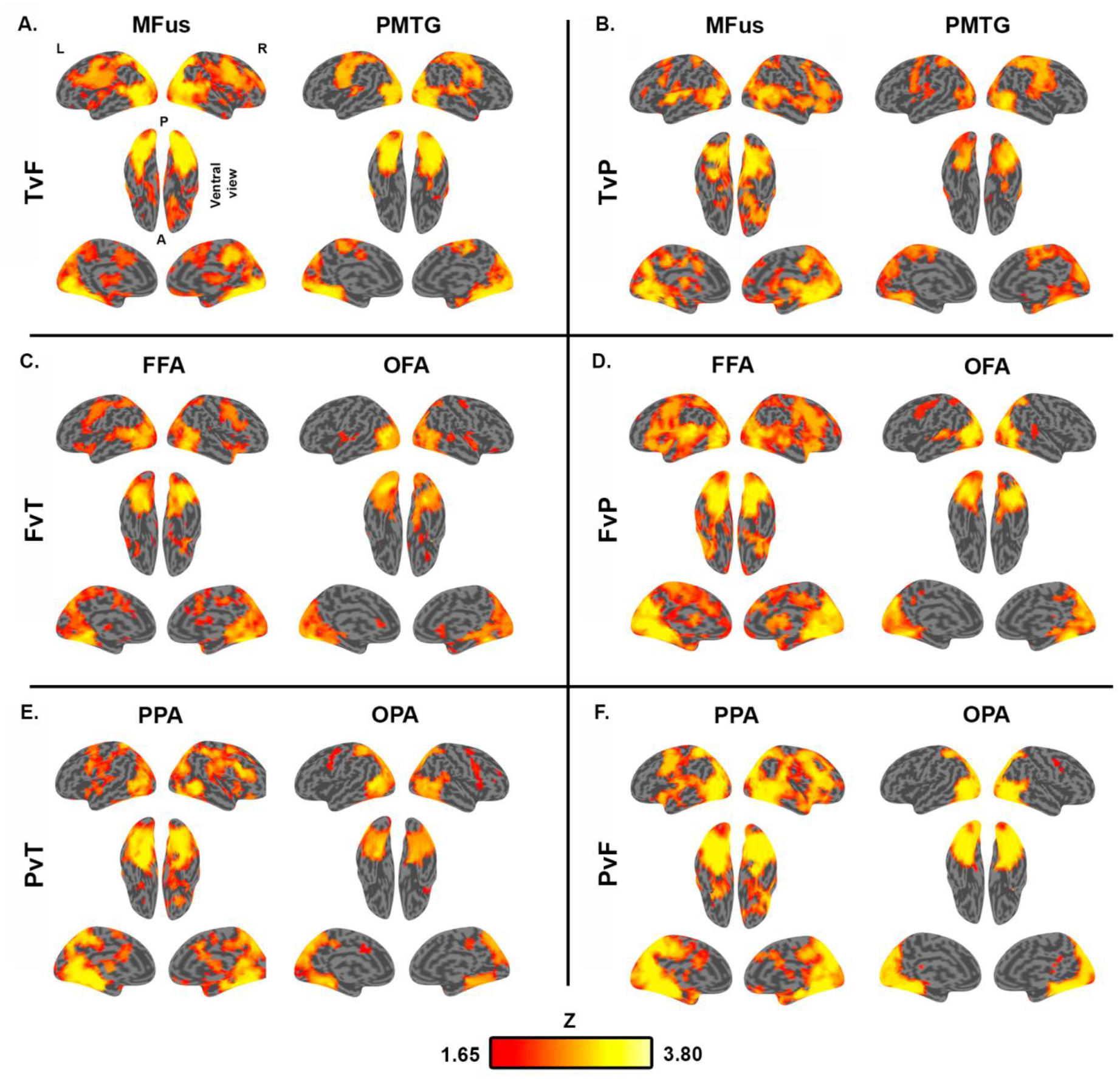
Group searchlight maps showing ‘good seed’ areas, for each target region. Z-score voxel intensities (threshold-free cluster enhancement paired t-test; Z-threshold > 1.65) show regions that seed significantly higher decoding accuracy for most-connected-than least-connected voxels within a given target region. Decoding comparisons: (a) tools vs. faces (TvF); (b) tools vs. places (TvP); (c) faces vs. tools (FvT); (d) faces vs. places (FvP); (e) places vs. tools (PvT); (f) places vs. faces (PvF). T ool regions: MFus = medial fusiform gyrus; PMTG = posterior middle temporal gyrus. Face regions: FFA = fusiform face area; OFA = occipital face area. Place regions: PPA = parahippocampal place area; OPA = occipital place area.

## Discussion

Here, we emphasize two main findings. First, complex functional responses in OTC are strongly related to patterns of connectivity to distal brain areas (i.e. grey matter voxel sets that share strong functional connectivity with the wider brain yield consistently better pattern discriminability than lesser-connected sets do, across all tested categories of information). These findings align with previous demonstrations that local OTC responses are shaped by distal connectivity with the wider brain (Amaral et al., under review; Chen et al., 2017; Garcea et al., 2019; Lee et al., 2019; Osher et al., 2016; Saygin et al., 2012; Saygin et al., 2016), and the more general proposal that functional brain responses are strongly determined by the integration of relevant information shared via structural and functional connectivity to the wider brain (Mahon & Caramazza, 2011; Osher et al., 2016; Park & Friston, 2013; Ruttorf et al., 2019; Saygin et al., 2012; Saygin et al., 2016; Sporns, 2004; Sporns & Zwi, 2004; Varela et al., 2001). Ultimately, local computations and the organization of representational content in OTC are dependent on interactions between connectivity-constrained neural assemblies that are likely dedicated to achieving particular computational goals (e.g. coordinated tool use, or face-to-face social interaction; Mahon, 2015; Op de Beeck et al., 2008; Peelen & Downing, 2017).

Second, most-connected voxels are not merely those that are most-activated, as shown by higher pattern discriminability for most-connected-relative to most-activated voxel sets in several regions (i.e. MFus, FFA, & OFA, and performed equivalently in all other regions), and further, the decoding advantage for most-than least-connected voxel sets remains when average activation (voxel t-values) of the two sets is constrained. These results are consistent with the observation that even voxels with weak amplitude responses may contribute meaningfully to pattern discrimination (e.g. Haxby et al., 2001; Kamitani & Tong, 2005; Weiner & Grill-Spector, 2010); as such, the ‘informativeness’ of weakly activated voxels may be captured via connectivity to the wider brain.

The decoding differences shown here between most-connected- and most-activated voxel sets might, at first glance, seem to conflict with previous evidence that emphasizes a statistical similarity between connectivity and activity measures; for example, category-specific activation in fusiform gyrus is correlated with the degree of functional connectivity to seed areas that share the same ‘category-preference’ (e.g. voxel-level activation to tool stimuli correlates with the voxel-level connectivity to ‘tool-preferring’ IPL; e.g. Chen et al., 2017; Amaral et al., under review). Similarly, while ‘matched-activation’ analyses shown here demonstrate a decoding advantage in most-connected-relative to least-connected voxel sets when controlling for average activation between the two sets, these analyses also show that connectivity and activity are certainly related (i.e. activation could not be matched between sets in all regions, for all subjects). We do not claim that that local activity and distal connectivity are completely independent (nor that they perfectly predict each other). Instead, we show that, when considering distributed functional responses, connectivity is a powerful means of identifying voxels that afford discriminability of high-level object representations. Thus, the present findings do not contradict previous work, but instead describe the relationship between connectivity and functional responses at a more complex level. Indeed, this is a valuable theoretical contribution given the widespread emphasis on distributed responses as a central functional organization principle of OTC and the wider brain (e.g. Haxby et al., 2001).

Importantly, what exactly might account for the representational differences – at the level of multivoxel patterns – between most-connected and most-activated voxel sets? By definition, most-activated sets sample voxels with the highest t-values, potentially sampling from closely-packed patches of voxels, whereas most-connected sets are comprised of a comparatively broader distribution of voxel responses. We speculate that these sets may differentially sample the heterogeneous functional responses of OTC. While ‘patchy’ organization of OTC is shown at a relatively coarse-grain (e.g. OTC is comprised of sparsely distributed and largely non-overlapping cortical patches that respond strongly to different types of information; Weiner & Grill-Spector, 2010), similar heterogeneous organization is also reflected at a finer spatial grain. For example, some voxel clusters within FFA respond preferentially to faces (compared to other objects), while other clusters show approximately equal tuning to multiple object categories (Çukur et al., 2013; Grill-Spector et al., 2006; Grill-Spector et al., 2007; Hanson & Schmidt, 2011); however, such responses may partially reflect responses to visual features that covary with certain categories rather than tuning to the categories themselves (e.g. Grill-Spector et al., 2006; Hanson & Schmidt, 2011; e.g. similar responses to faces and round-shaped objects, such as clocks or apples, for both voxels and single-cell recordings in macaque inferior temporal cortex; Moeller et al., 2017; Tsao et al., 2006). As such, distributed cortical representations are comprised of heterogeneous voxel responses that reflect sensitivity to a diverse array of visual or semantic features, and such sparse encoding may allow for an exhaustive representational capacity of OTC via complex response patterns (Grill-Spector et al., 2006; Olshausen & Field, 2004).

Accordingly, we suggest that most-connected voxel sets may, in some cases, advantageously sample relatively more-diverse information than most-activated voxel sets. At a cognitive level, the connectivity-based voxel selection approach may better-exploit computations occurring within heterogeneous patches dedicated to different types of domain-specific information. For instance, subsets of FFA voxels with strong connectivity to OFA may reflect greater tuning to face-parts, while voxels that are well-connected to STS may be preferentially tuned to dynamic-emotion related information – potentially indexing the integration of information between different patches within a domain-specific network. Thus, voxel selection by connectivity recruits voxels that are highly connected with distal areas, bringing about a diverse set of object-related information. Contrastingly, selection by local activity targets voxels with strong amplitude responses that are potentially very important for the particular computations at play within that region. At the neural level, the connectivity-based approach may sample widely from functionally discrete patches, while the activity-based approach may draw more from spatially clustered set of voxels with very similar (i.e., less informationally diverse) response profiles (Bell et al., 2009; Çukur et al., 2013; Grill-Spector et al., 2006; Grill-Spector et al., 2007).

For example, given 5 functionally discrete patches within a given target area, most-connected voxel sets may be more likely to sample from each patch than most-activated sets that may draw more heavily (and potentially, more redundantly) from fewer patches that exhibit strong, clustered amplitude responses. Thus, most-activated voxel sets may, in some cases, suffer from a higher degree of ‘informational redundancy’.

In the current study, we demonstrate higher decoding in most-connected-than least-connected voxel sets when using a-priori seed regions (e.g. tool-preferring PMTG, IPL, & SPL, were used as seeds for voxel selection and tool decoding in MFus), as motivated by highly correlated resting-state activity between areas that share category preferences (Kamps et al., 2020; Stevens et al., 2015; Zhang et al., 2009; Zhu et al., 2011). However, searchlight analyses revealed that regions outside of these designated areas also afford similar effects, perhaps with some of these connections sub-serving both bottom-up and top-down modulations of local signal. These results are consistent with previous research showing that OTC sub-regions (e.g. FFA) show strong functional connectivity to regions that sub-serve more domain general (and task-relevant) processing, that are often considered key nodes (e.g. posterior parietal cortex and inferior frontal gyrus) among attention- or cognitive control networks (Cole et al., 2010; Hutchinson et al., 2014; Vogel et al., 2012).

We note that our central claim in this paper (i.e. that local computations are influenced by connectivity to – and presumably via computations within – distal brain areas) is *directionally agnostic*; that is, from the present data, we cannot claim that local computations are causally influenced by connectivity to distal brain areas (or vice versa). Instead, future research may address the causal nature of this relationship with neural disruption measures (e.g. transcranial magnetic stimulation) or brain lesion studies. We also note that the connectivity-based voxel selection approach used here is potentially generalizable to most other fMRI decoding experiments.

In conclusion, the current data shows that high-level multivariate representations in OTC can be reliably indexed by functional connectivity, demonstrating the importance of connectivity constraints on the functional organization of OTC.

## Acknowledgments

Special thanks to B.Z. Mahon, P.E. Downing, & K. Koldewyn for helpful discussions with the manuscript; D. Lee, S. Kristensen, L. Amaral & F. Bergstrom for data collection.

## Author Contributions

JW designed the experiment, analysed data, and wrote the manuscript; JA designed the experiment and analyses, and wrote the manuscript.

## Data availability and code availability statement

The data and accompanying code is available upon request from the authors.

